# A simple molecular mechanism explains multiple patterns of cell-size regulation

**DOI:** 10.1101/083725

**Authors:** Morgan Delarue, Daniel Weissman, Oskar Hallatschek

## Abstract

Increasingly accurate and massive data have recently shed light on the fundamental question of how cells maintain a stable size trajectory as they progress through the cell cycle. Microbes seem to use strategies ranging from a pure sizer, where the end of a given phase is triggered when the cell reaches a critical size, to pure adder, where the cell adds a constant size during a phase. Yet the biological origins of the observed spectrum of behavior remain elusive. We analyze a molecular size-control mechanism, based on experimental data from the yeast *S. cerevisiae*, that gives rise to behaviors smoothly interpolating between adder and sizer. The size-control is obtained from the titration of a repressor protein by an activator protein that accumulates more rapidly with increasing cell size. Strikingly, the size-control is composed of two different regimes: for small initial cell size, the size-control is a sizer, whereas for larger initial cell size, is is an imperfect adder. Our model thus indicates that the adder and critical size behaviors may just be different dynamical regimes of a single simple biophysical mechanism.

---

Cells need to coordinate growth and division to keep their size in the physiologically optimal range (1–4). A range of mechanistic models for how cells can link division to growth have been proposed (5–10), and painstaking experimental work has uncovered much of the network of genes and reactions involved (3, 4, 11, 12). Recently, high-throughput experimental techniques have enabled the detailed measurement of size dynamics at the level of single cells (13–16), revealing in detail how cells fluctuate around their typical size; these results have been described by phenomenological models that focus purely on the size dynamics (16, 17). Here we connect the two approaches by exploring the “titration” model of cells’ response to size fluctuations (5, 7, 9, 18) (also known variously as the “concentration”, “inhibitor-dilution”, or “structural” model depending on the interpretation (7)) and showing that it can give rise to the observed phenomenological patterns.

The basic idea of the titration model is that a transition is triggered when the concentration of an activator protein exceeds that of a repressor or inhibitor. There are two key steps needed to connect this model to the observed single-cell size data. First, while the simplest versions of the model assume that the activator concentration is initially negligible and the amount of repressor is constant (see (7), “structural model”), we consider potential ways in which they might vary across cells and over time. Second, while many phenomenological models focus on describing the total change in size over one whole cell cycle (7, 17), cells regulate their size only in certain phases of the cell cycle, such as the B and D intervals in *E. coli* (19), or the G1 or G2 phases of the cell cycle for the budding yeast *S. cerevisiae* (20). Thus, we use the titration model to describe the regulation of a single phase of the cell cycle, with the full size regulation being composed of a series of such steps involving different pathways.

We focus on the first phase of the budding yeast *S. cerevisiae* cell cycle, the G1 phase (from birth to bud), for which recent experiments provide detailed information on regulation at the single-cell and molecular levels (20–22). In this case, the activator and repressor are, respectively Cln3 and Whi5, which react in the cell nucleus (23, 24). The key to their usefulness in sensing cell size are their different patterns of production and degradation. Cln3 molecules are produced in the cytosol during G1 at a rate proportional to cell volume (22) and degraded at a (fairly rapid) constant rate (25), while Whi5 is neither produced nor effectively degraded during G1 (22), and so remains constant in number. Thus, as the cell grows, the number of Cln3 molecules eventually exceeds the number of Whi5 molecules, and the cell exits G1. Mathematically, if *v*(*t*) is the cell volume at time *t* after birth, *κ*_*p*_ is the rate density at which Cln3 is produced per unit volume, and *k*_*d*_ is the rate at which it degrades, the number of Cln3 molecules *N*_*a*_(*t*) evolves as

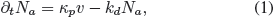

with the end of G1 being triggered when *N*_*a*_ > *N*_*r*_, the (constant) number of Whi5 molecules.

We wish to analyze the predictions of this model for the dynamics of cells that are born with varying initial sizes *v*_0_. To do this, we need to understand the boundary conditions of Eq. (1). The initial concentration *c*_*a*,0_ of Cln3 is roughly constant (22), consistent with it being close to an equilibrium of Eq. (1) outside of G1. In contrast, the absolute number of Whi5 is roughly constant, independent of initial cell size, as it is produced at a size-independent rate (22) during the budded phase, whose duration is not dependent on cell size. <u>How</u> the cell synthesizes Whi5 at a constant rate independent of its size is an interesting question that we will not attempt to answer here, beyond noting that one possibility is that transcription, rather than translation, may be the rate-limiting step. In the Supplementary Information, we show that this differing dependence of the number of activator and repressor molecules on the volume at birth is crucial to maintaining size control. Note that the differing dependence implies that for extremely large cells, there will already be sufficient activator at birth; at this point, the model breaks down and other reactions that are normally rapid compared to the duration of G1 will set the timescale. This breakdown of the model can explain why mother budding yeast cells, which keep on increasing in birth size generations after generations, do not seem to exhibit any size-control in G1 (26).

As the cell volume monotonously increases with time, we can replace time in Eq. (1) with cell volume: ∂_*t*_• = ∂_*t*_*v* × ∂_*v*_•. Budding yeast grows exponentially during the G1 phase (21), i.e., ∂_*t*_*v* = *kv* for some growth rate *k*. (In the Supplementary Information, we also consider a linear growth model, which gives qualitatively similar results.) Eq. (1) thus becomes:

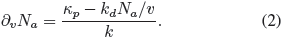

The volume of the cell increases until the activator matches the repressor upon which the next phase of the cell cycle is entered. Solving *N*_*a*_(*v*_*f*_) =*N*_*r*_ gives the final volume *v*_*f*_ at the end of G1. The exact solution is complicated (Supplementary Information), but exhibits two simple limiting regimes (Figure 1A):

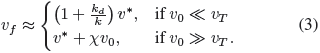

**Figure 1:**
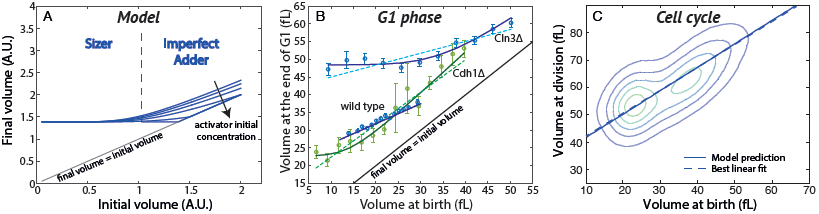
Model and fit of experimental data. A. The model predicts two regimes of size control: a “critical sizer” for small cells and an “imperfect adder” for large cells. B. Volume at the end of G1 in the yeast *S. cerevisiae*, for different mutants, as a function of the volume at birth. Data from (20) (unpublished). Each point shows the mean ± SEM. The lines correspond to the best fit from our model. Dashed lines correspond to best linear fits. C. Volume at division as a function of volume at birth in yeast. The line shows the prediction for our model, and the dashed line shows the linear fit. Data from (20).

Here *v*\* = *kN*_*r*_/*κ*_*p*_ is the volume scale determined by the basic titration mechanism, χ = (1 − (*k* + *k_d_*)*c*_*a*,0_/*κ_p_*)^*k*/(*k_d_*+*k*)^ gives the strength of the dependence of *v*_*f*_ on *v*_0_, and threshold volume separating the regimes is *v*_*T*_ = *v*\**k*_*d*_/(*k*_χ_).

Eq. (3) says that for cells that are born small, *v*_0_ ≪ *v*_*T*_, almost all of the activator is produced near the end of G1, and the mechanism enforces a minimum final volume independent of the volume at birth, i.e. a “critical size”, as described in (27) for the fission yeast. For cells that are born large, *v*_0_ ≫ *v*_*T*_, the activator is produced more evenly throughout G1 and the initial number may also be important, and the final volume is an affine function of the volume at birth. In the limit where the initial activator concentration is small, as in the simple concentration model (7, 28), the slope χ = 1 and the mechanism acts as an “adder” (the “incremental” model, (17)). More generally, for χ ≠ 1, the mechanism acts as an “imperfect adder” (16). Note that the threshold size *v*_*T*_ may be very different from the typical size of cells, so that almost all cells in a population may exhibit the same phenomenological pattern of size control (e.g., almost all cells may be “large” in this sense).

To test our predictions, we reanalyzed recent data on the budding yeast *S. cerevisiae* from Soifer and colleagues ((20), unpublished data). In their study, they systematically measured the volume of cells at the beginning and end of G1, for 520 single-gene knock-out mutants affecting cell size. For the wild type, we observe a single affine regime with a slope χ = 0.56, indicating an imperfect adder (Figure 1B). 490 of the 520 mutants are also mostly in the imperfect adder regime. The distribution of slopes is peaked around the wild-type value but broadly distributed, with standard deviation σ(χ) = 0.2 (Supplementary Figure S3A), indicating that χ is not robust to mutation. The remaining 30 mutants cannot be described by a simple affine relationship between *v*_0_ and *v*_*f*_ (e.g., Figure 1B, Cln3 and Cdh1 mutants; see Supplementary Information for all 30 mutants). For all 30, we find exactly the pattern predicted by the model: a critical sizer for small cells crossing over to an imperfect adder for larger ones. Note that the fact that the Cln3 knockout still controls its size indicates the existence of compensatory mechanisms, and the accumulation of another activator, possibly Cln1 or Cln2.

Finally, we consider recent experiments by I. Soifer and colleagues showing that budding yeast are adders over their whole cell cycle (29), even though the size-control in G1 is an imperfect adder (20). We find that our model can can give rise to an adder *over the whole cell cycle* by considering that different phases of the cell cycle have different size-control mechanisms. In contrast to G1, the phase between the end of G1 and division is a “timer” (20, 22): its duration, of about *t*\* = 50 minutes, is largely independent of cell size at the end of G1. In this case, the volume at division *v*_*d*_ is simply proportional to the volume at the end of G1 *v*_*G*1_:

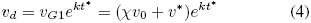

Using the independently-measured parameter values *t*\* = 50 minutes, *k* = 0.0096 min^−1^ (21), Eq. (4) is in excellent agreement with the experimental data (29) (Figure 1C). The fact that two independent size control mechanisms, the imperfect adder in G1 and the timer following G1, have been tuned to combine to give a near-perfect adder (χ*e^kt*^* = 0.92) suggests that there has been selection for the adder phenotype at the whole cell-cycle level. Soifer et al show that the adder phenotype could be a caused by cells producing a fixed amount of Whi5 in the budded phase (29). Their model is similar to ours, except that it predicts that the amount of Whi5 varies with cell size at birth (see Supplementary Information), and neglects the dynamics of Cln3 concentration, leading it to predict just a single regime of size regulation in G1.

In this letter, we analyze the titration model of size control, in which a phase of the cell cycle ends when the concentration of an activator protein exceeds a threshold set by a repressor protein. We find that for size control to be maintained, not only the activator and repressor need to accumulate differently as the cell’s volume changes during the phase in question, but they also need to respond differently to volume changes <u>before</u> the phase begins. Given that these conditions are met, we predict that any single genotype will show two different patterns of size control, depending on the initial cell volume: a critical size for small initial volumes, and an imperfect adder for large initial volumes. We find this two-stage pattern in many mutant strains of yeast. All other observed strains are consistent with the model, in that they follow the imperfect adder pattern. We predict that for these strains, careful measurement of rare small cells would reveal the critical sizer regime; such measurements may be practical with microfluidic techniques (30). These techniques could also enable the observation of the anticipated breakdown of the model for rare extremely large cells.

While we have focused on budding yeast, the model is generic and does not rely on the molecular details of the full cell cycle pathway; even in yeast, the activator and repressor in the model may be effective quantities that do not correspond exactly to Cln3 or Whi5. Indeed, nothing in the model requires that the “repressor protein” even be a protein; it could just as well be any set of physical sites whose number does not scale linearly with cell volume (7), such as genomic regions which need to be bound by transcription factors to trigger DNA replication, or locations on the cell membrane to which the activator need to bind to trigger division. Thus, we predict that the same patterns of size regulation will also be found in other species. For example, while fission yeast *Schizosaccharomyces pombe* acts as a critical size for typical initial cell volumes (31), high-throughput microfluidics experiments have enabled the measurement of size dynamics in the tails of the distribution (15), revealing the predicted second regime (Supplementary information). The bacterium *E. coli* may also be an example, as our model is in better agreement with observed phenomenological size dynamics (14) than a simple imperfect adder scenario (Supplementary Information), although again more observations of unusually small cells are needed to make a definitive statement.

## ACKNOWLEDGMENTS

The authors would like to thank I. Soifer for kindly providing additional experimental data.

